# TDP-43 pathology induces CD8^+^ T cell activation through cryptic epitope recognition

**DOI:** 10.1101/2025.06.22.660773

**Authors:** Shahab Chizari, Matteo Zanovello, Steven Kong, Vidur Saigal, Anna-Leigh Brown, Valentina Turchetti, Bingxian Chen, Connor C. Devine, Luca Zampedri, Iwona Skorupinska, Giacomo Maria Minicuci, Francesca Paron, Paola Tonin, Giulia Marchetto, Ziyi Li, Jennifer M. Colón-Mercado, Simone Barattucci, Darija Soltic, Dario Dattilo, Ariana Gatt, Jada M. Hembrador, Guido Capasso, Federica Frezzato, Livio Trentin, Elisabeth Lin, Raquel Lopes, Nathan Routledge, Yue A. Qi, Michael G. Hanna, Michael Ward, Leonard Petrucelli, Maurizio Romano, Gaetano Vattemi, Emanuele Buratti, Andrea Malaspina, Ashirwad Merve, Pedro M. Machado, Gianni Sorarù, Pietro Fratta, Ning Jiang

## Abstract

Aggregation and nuclear depletion of the RNA binding protein TDP-43 are the crucial pathological features of amyotrophic lateral sclerosis (ALS) and inclusion body myositis (IBM), two degenerative diseases of the CNS and muscle. The loss of TDP-43 nuclear function results in the aberrant inclusion of cryptic exons in mRNA transcripts, leading to the expression of *de novo* proteins. Clonally expanded and highly differentiated CD8^+^ T cells have been observed in individuals with TDP-43 proteinopathies and therapeutics modulating the T cell response have recently been found to extend survival. However, the target antigens mediating T cell activation have remained elusive. Here, we investigate whether the *de novo* proteins induced by aberrant cryptic splicing due to TDP-43 nuclear loss can act as neo-antigens. We detect the HDGFL2 cryptic peptide and multiple other TDP-43 cryptic exons in IBM skeletal muscle, where their presence correlates with enrichment of T cells and class I antigen presentation pathways. Furthermore, we identify epitopes deriving from HDGFL2 and IGLON5 cryptic peptides which are recognized by clonally expanded and functionally differentiated populations of CD8^+^ T cells in ALS and IBM Patients. Finally, we demonstrate that T cells engineered to express the identified TCRs can bind and activate in response to the cryptic peptide derived epitopes (cryptic epitopes) and are able to kill TDP-43 deficient astrocytes. This work identifies for the first time specific T cell antigens in ALS and IBM, directly linking adaptive immune response to TDP-43 pathology.

## Introduction

Aggregation and nuclear depletion of the RNA-binding protein TDP-43 is a defining pathological feature of amyotrophic lateral sclerosis (ALS), inclusion body myositis (IBM), and several other neurodegenerative conditions^1,2^. TDP-43 is a ubiquitously expressed protein that resides predominantly in the nucleus, where it has been shown to be a repressor of cryptic exons, intronic sequences that are normally excluded from mRNA^3^. Upon TDP-43 mislocalisation to the cytoplasm, these cryptic exons become derepressed and incorporated into transcripts. Although most of these events generate frameshifts and premature stop codons leading to RNA degradation^4^, we and others have shown that some cryptic exons can be translated to form cryptic peptides in the brain of patients with ALS^5,6^. Similarly, loss of TDP-43 mediated splicing repression also generates cryptic exons and cryptic peptides in IBM skeletal muscle^7,8^.

Highly differentiated and clonally expanded CD8^+^ T cell populations have recently been described to be enriched in ALS^9,10^ and IBM^11^. These cell populations exist both in the peripheral blood mononuclear cells (PBMCs)^9^ and in disease relevant tissue such as cerebrospinal fluid (CSF)^10^ and skeletal muscle^11^. T cell clonal expansion in addition to the increased expression of proteins associated with activation and cytotoxicity supports an antigen-specific T cell response^12^ however, no clear antigenic targets have been described, and attempts at finding them have largely been unsuccessful^13–15^. CD8^+^ T cells can recognize novel peptides derived from non-canonical reading frame translation in cancer cells^16,17^, raising the crucial question of whether CD8^+^ T cells can mount a response against cells expressing cryptic epitopes in TDP-43 proteinopathies.

Here, we examined IBM skeletal muscle using immunohistochemistry, proteomics, and RNA sequencing to further evaluate the cryptic exon expression pattern and signature for immune activation (Fig. 1a, left). We detected the HDGFL2 cryptic peptide co-localized with TDP-43 loss along with widespread cryptic exon expression which correlated with T cell infiltration and class I antigen presentation pathways in the diseased tissue. We then utilized a high-throughput and multidimensional single T cell profiling approach, TetTCR-SeqHD^18,19^, to show that cryptic epitope specific CD8^+^ T cells isolated from ALS and IBM PBMCs are polyclonal, highly clonally expanded, highly differentiated and functional, and capable of binding multiple cryptic epitopes, including those derived from HDGFL2 and IgLON5 (Fig. 1a, right). Additionally, we showed that cells engineered to express HDGFL2 and IgLON5 cryptic epitope specific T cell receptors (cryptic-specific TCRs) are capable of not only binding the cryptic epitopes, but also mediating T cell specific activation and effector responses. Furthermore, we demonstrated that TDP-43 knockdown (KD) in an astrocyte cell line led to HDGFL2 cryptic exon expression as well as activation and cytotoxicity of primary CD8^+^ T cells transduced to express an HDGFL2 cryptic-specific TCR. This work highlights for the first time T cell antigenic targets in ALS and IBM, linking them to the crucial pathological hallmark of these disorders.

**Fig. 1.**
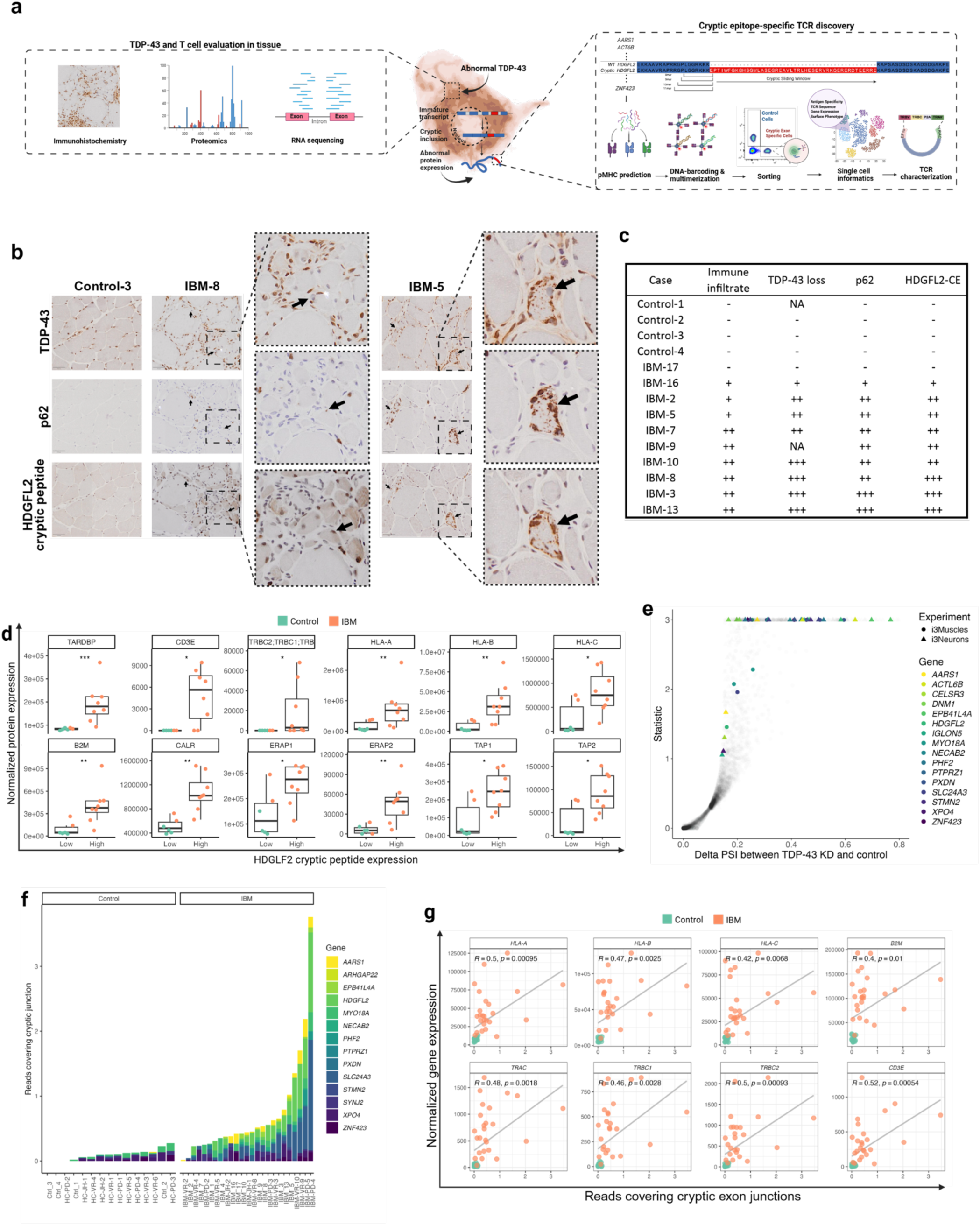
Cryptic peptides and MHC I / TCR pathways are found in IBM tissues. **(a)** Schematic of the experimental workflow. IHC, proteomics, and RNA sequencing were used to analyze the expression of cryptic peptide, TCR, and MHC expression and T cell infiltration in IBM muscle tissues (left). Validated cryptic peptides were used to informatically predict epitopes binding to patient specific HLA molecules. Then, fluorescence labeled and DNA-barcoded pMHC tetramers were generated and used to sort cryptic epitope Tetramer^+^ CD8^+^ T cells for downstream single cell analysis and *in vitro* characterization (right). **(b)** Visualization of seriate immunohistochemistry staining for TDP-43, p62, and HDGFL2 cryptic peptide in muscle from one control and two IBM cases. For each case, arrows show colocalisation of HDGFL2 cryptic peptide in fibers with p62 and TDP-43 nuclear depletion and/or aggregation. **(c)** Semi-quantitative scores for immune infiltrates, TDP-43 loss, p62 and HDGFL2 cryptic peptide in control and IBM cases. **(d)** Normalized protein expression as quantified by mass spectrometry. The same group of four controls and 10 IBM cases were grouped based on the expression level of HDGFL2 cryptic peptide quantified by antibody in (c). Box plots show median and all points are shown. **(e)** Volcano plot showing upregulated splicing junctions upon TDP-43 knockdown in iPSC-derived skeletal muscles (circles) and iPSC-derived glutamatergic cortical neurons (triangles). **(f)** Normalized counts of RNA-seq reads covering cryptic exon junctions in controls (n = 15) and IBM cases (n = 25). **(g)** Correlation between RNA-seq normalized read counts for TCR ɑ and β constant chain genes (*TRAC*, *TRBC1*, and *TRBC2*), *CD3E*, *HLA-A*, *HLA-B*, *HLA-C*, and β2-microglobulin (*B2M*) and cryptic exon normalized counts in controls (n = 15) and IBM cases (n = 25).

### Cryptic peptides are present in IBM skeletal muscle and correlate with TCR and MHC-I gene expression

TDP-43 aggregation and T cell infiltration in skeletal muscle are hallmarks of IBM^11^, and recent work has suggested that degenerative features such as rimmed vacuoles, TDP-43 loss of function, and cryptic exons expression are present in IBM tissue^7^. Recently, two different groups have developed antibodies that bind to the HDGFL2 cryptic peptide^6,20^. Using these antibodies, they showed that the HDGFL2 cryptic peptide can be detected in CSF and blood of presymptomatic and early-stage of ALS/FTD individuals^6^ and that it is significantly increased in brain regions with TDP-43 pathology in frontotemporal lobar degeneration with TDP-43 pathology (FTLD-TDP)^20^ and Alzheimer’s disease with TDP-43 pathology (AD-TDP)^20^, as well as in muscle biopsies from IBM cases affected by TDP-43 pathology^8^, compared to non-TDP-43 controls.

To further investigate HDGFL2 cryptic peptide in IBM patient muscle, we performed immunohistochemical staining (IHC) for HDGFL2 cryptic peptide using a newly generated antibody^20^, TDP-43, and p62 (an autophagy component that colocalizes with protein aggregates), in 10 IBM patients and four controls (Supplementary Table 1). We were able to visualize the specific accumulation of TDP-43 in p62-positive aggregates, with co-occurrence of TDP-43 nuclear loss in some fibers (Fig. 1b,c). Importantly, we were able to detect HDGFL2 cryptic peptide expression in 9/10 IBM cases and 0/4 controls (Fig. 1b,c), and we found immune infiltrates in samples with higher levels of TDP-43 aggregation (Fig. 1b,c). Intriguingly, HDGFL2 cryptic peptide showed a similar localisation to TDP-43 and p62, suggesting it may also aggregate in affected IBM myofibers.

We then used proteomics data to evaluate the relationship between the presence of the HDGFL2 cryptic peptide with antigen processing/presentation and T cell infiltration. Strikingly, CD3ε and TCRβ proteins were virtually absent in controls and expressed only in a subset of IBM cases with higher expression level of the HDGFL2 cryptic peptide (Fig. 1d). Proteins in the MHC-I peptide processing and presentation pathway and HLA proteins were also significantly higher in these cases (Fig. 1d and Extended Data Fig. 1a). Differential expression and GO enrichment analysis showed a reduction of muscle-specific terms and increased antigen processing and presentation terms in the IBM cases compared to controls (Extended Data Fig. 1b,c).

As TDP-43 depletion leads to the expression of many cryptic peptides (Fig. 1e), we aimed to assess the broader relationship between TDP-43 loss of function and T cell responses in IBM. We performed RNA sequencing in TDP-43 deficient iPSC-derived myocytes to identify high-confidence muscle CE events predicted to give rise to cryptic peptides. We then investigated their presence in patient tissue using RNA sequencing data from 25 IBM muscle tissues and 15 controls (Extended Data Fig 1d,e and Extended Data Fig. 2b)^7^. We detected multiple cryptic exons in the muscles of IBM patients, while being almost undetectable in controls (Fig. 1f). *HDGFL2* cryptic exon showed the highest specificity (0/15 healthy controls) and sensitivity (17/25 IBM cases). A number of cryptic exons such as *ACTL6B* and *IgLON5*, which are found in brain tissue from people affected from ALS (Supplementary Data 2), were not identified in the IBM cohort, possibly due to the low expression of these genes in muscles (Extended Data Fig. 2c)^21^. Intriguingly, we detected the *STMN2* cryptic exon, which has neuronal specificity^22^, in some IBM cases (Extended Data Fig. 2d). To address whether TDP-43 loss of function and the T cell response are correlated in diseased tissue, we used the cumulative detection of cryptic exons as a measure of TDP-43 loss of function burden, and tested its relation to the expression of antigen presentation (MHC-I) and T cell infiltrate (TCR and CD3 transcripts). We found an elevated expression of TCR and HLA genes in the IBM cases (Extended Data Fig. 2e), and we observed a strong correlation between cryptic exon burden and the expression of both TCR and HLA genes indicating a link between these events in disease (Fig. 1g).

### Cryptic epitope Tetramer^+^ T cells are enriched and clonally expanded in the PBMCs of ALS and IBM

To address whether CD8^+^ T cells are involved in the response to cryptic peptide in ALS and IBM, we used our previously established DNA-barcoded pMHC tetramer-based high-throughput multiomic single cell profiling technology, TetTCR-SeqHD^18,19^ (Fig. 1a, right). From our RNA sequencing of IBM muscle and TDP-43 deficient iPSC-derived muscles and neurons along with previously validated cryptic exons from neuronal cells^5^, we defined a complete list of cryptic peptide sequences as T cell epitope candidates (Fig. 1e). We employed a suite of pMHC binding algorithms^23^ to computationally predicted a panel of 379 cryptic epitopes for CD8+ T cells, covering 18 cryptic peptides and 9 HLA-alleles, to generate DNA barcoded pMHC tetramers (Tetramer).

CD8^+^ T cells enriched from the PBMCs from ALS (N=9) and IBM (N=4) patients, and healthy controls (N=7) were stained with a donor-specific cryptic epitope Tetramer panel, a control viral Tetramer panel, along with a panel of fluorophore labeled and DNA-barcoded phenotyping/sample-tagging antibodies. Tetramer^+^ cells were sorted for high-throughput amplification of T cell-bound Tetramer DNA-barcodes TCR genes, as well as 558 genes and 17 surface proteins indicative of T cell activation and differentiation. There were significantly higher cryptic epitope Tetramer^+^ (CE-Tetramer^+^) cells in ALS and IBM patients compared to healthy controls (Fig. 2a,b). Importantly, in ALS and IBM cases, CE-Tetramer^+^ CD8^+^ T cells were significantly more clonally expanded (Fig. 2c), with merely 3.7% of CE-Tetramer^+^ T cells expanded in healthy donors (>2 cells in a clonotype), compared to the 31% and 37.8% in ALS and IBM donors respectively. Furthermore, “hyperexpanded” (>15 cells with the same TCR sequences) and “large” (>10 cells with the same TCR sequences) CE-Tetramer^+^ T cell clones were only identified in IBM and ALS donors (Fig. 2d).

**Fig. 2.**
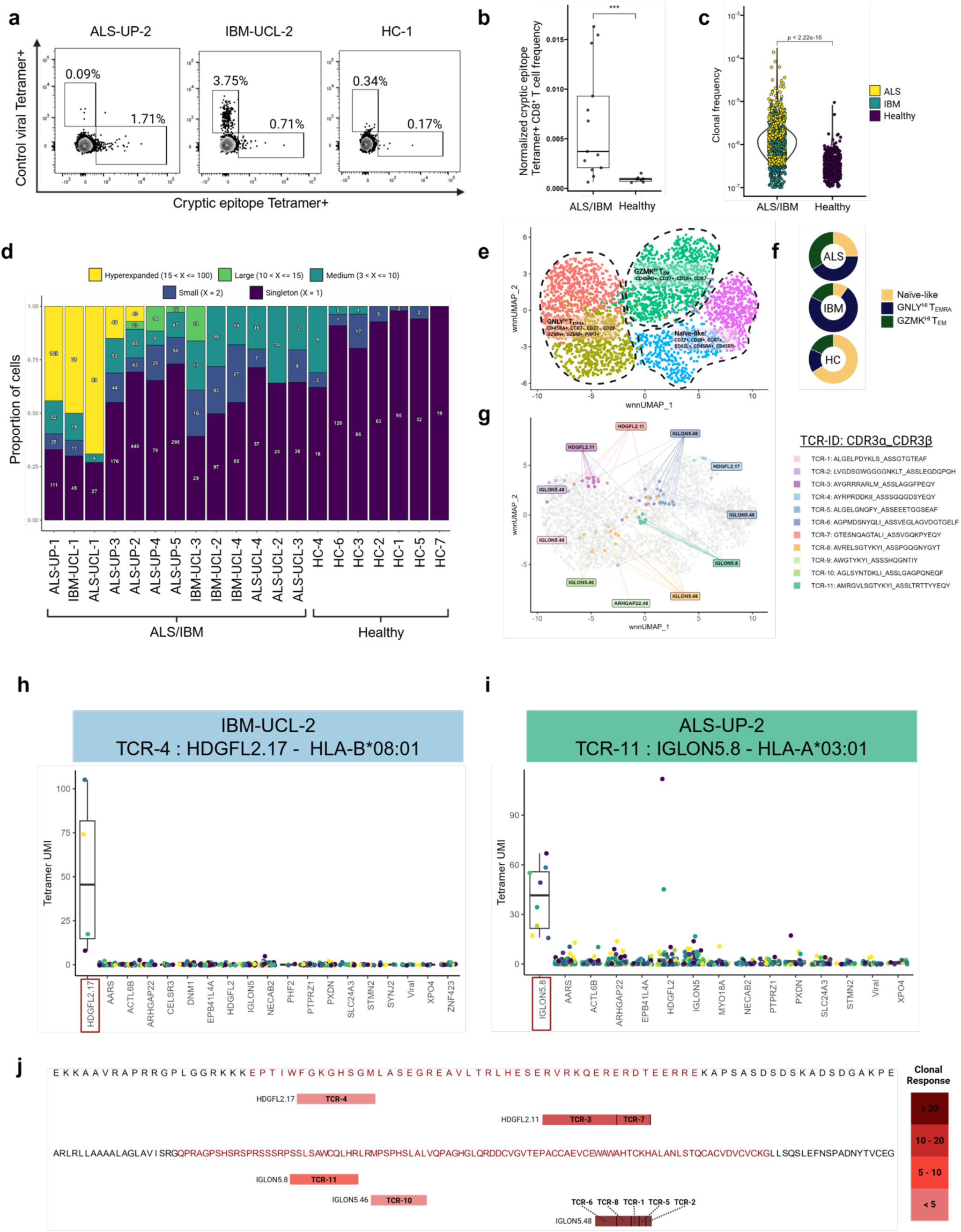
TDP-43 dysfunction leads to a heterogenous and polyclonal CD8^+^ T cell response to cryptic epitopes. **(a)** Representative FACS plot depicting cryptic epitope Tetramer^+^ cells (x-axis) and control viral Tetramer^+^ cells (y-axis) across three donors. **(b)** Summary of the complete data represented in (A). Cryptic epitope Tetramer^+^ CD8^+^ T cell frequency was normalized by the number of total pMHC Tetramers for each individual donor. Wilcoxon Rank Sum test was used to evaluate significance. **(c)** Violin plot of clone size normalized by starting number of CD8**^+^** T cells across ALS, IBM, and healthy donors. The Wilcoxon Rank Sum test was used to evaluate significance. **(d)** T cell clonality per donor ordered by greatest to least clonality. Clones were stratified by their CDR3β according to the following bins: Hyperexpanded (15 < X <= 100), Large (10 < X <= 15), Medium (3 < X <= 10), Small (X = 2), Singleton (X = 1). **(e)** UMAP representation of single cells by a weighted nearest neighbor calculation of gene and surface protein expression. Clusters were manually annotated based on their differential gene and protein expression. Genes are italicized. **(f)** Donut graph of phenotypic composition across ALS, IBM, and healthy controls. **(g)** Tetramer specificity assignments on T cell clones overlaid on UMAP. Clonal specificities were assigned such that at least 60% of the cells in a given clonotype must share the same epitope assignment based on tetramer barcode sequencing. Cells that do not meet that threshold are colored grey. **(h/i)** Box plot visualization of tetramer barcode UMI counts for TCR-4 and TCR-11 with assigned specificities to HDGFL2.17 (left) and IgLON5.8 (right) epitopes. Each dot depicts an individual cell’s signal for each tetramer barcode (x-axis) within the specified clone. All tetramer barcodes were grouped into their parent cryptic protein except for the top hit (represented by the red box). All points are shown and center line represents median **(j)** Depiction of identified cryptic epitopes across HDGFL2 and IgLON5. Epitopes found to be targeted are marked by a bar which spans the epitope sequence. The tick marks within each bar represent different T cell clones targeting the same epitope and the distance within each tick represents the relative clone size. The color intensity represents the sum of all T cell clones targeting the particular epitope.

### CE-Tetramer^+^ T cells are associated with activation and cytotoxicity phenotypes

To gain more information on the subtype and status of the identified CE-Tetramer^+^ T cells, we performed weighted-nearest neighbor (WNN) analysis and subsequent UMAP visualization of our single cell gene and surface protein expression data (Fig. 2e). We observed five clusters across three major phenotype groups including: Naive-like (CD27^+^, CD28^+^, CCR7^+^, CD62L^+^, CD45RA^+^), GZMK^HI^ T effector memory (T_EM_) (CD27^+^, CD28^+^, CCR7^-^, CD45RO^+^, GZMK^+^), and GNLY^HI^ T effector memory RA (T_EMRA_) (CD27^-^, CD28^-^, CCR7^-^, CD45RA^+^, GNLY^+^). CE-Tetramer^+^ CD8^+^ T cells from ALS and IBM patients were enriched for the GZMK^HI^ T_EM_ and GNLY^HI^ T_EMRA_ populations, whereas CE-Tetramer^+^ CD8^+^ T cells from healthy control were largely Naive-like (Fig. 2e,f). Previous single cell RNA sequencing of CSF T cells isolated from AD and Parkinson’s disease (PD) patients show a similarly high expression of related cytolytic and granzyme genes associated with a T_EMRA_ phenotype^24,25^.

### CD8^+^ T cells are capable of mounting polyclonal responses to HDGLF2 and IGLON5 cryptic epitopes

We combined single cell Tetramer sequencing data with TCR sequencing data to assign epitope specificities to T cell clones. To generate high confidence assignments, T cell clones (>3 cells in a clone) were given a particular cryptic epitope specificity only when at least 60% of all cells within the TCR clonotype share the same maximum tetramer barcode signal. Under these conditions, 11 T cell clones were assigned specificities, 10/11 of which recognized cryptic epitopes derived from either HDGFL2 or IGLON5 (Fig 2g-j). These clones were found across the GZMK^HI^ T_EM_ and GNLY^HI^ T_EMRA_ clusters (Fig 2g).

To experimentally validate the specificity and activation ability of the above-identified cryptic epitope specific (cryptic-specific) TCRs, we engineered J76-CD8 T cells, a human CD4^+^ T cell line engineered with a human CD8 molecule, to express the clonally expanded TCRs that we found bind to the epitopes derived from HDGFL2 and IgLON5 through our high-throughput screen.

We found that two TCRs were able to robustly bind their cognate cryptic epitope and become activated, one originally identified from an IBM patient, TCR-4, and the other from an ALS patient, TCR-11 (Fig. 3a). TCR-4, derived from donor IBM-UCL-2, was able to bind a cryptic epitope sequence from HDGFL2 cryptic peptide when presented on HLA-B*08:01. TCR-11, derived from donor ALS-UP-2, was able to bind a cryptic epitope sequence from IgLON5 cryptic peptide when presented on HLA-A*03:01. We then tested the ability of these TCRs to induce T cell activation by co-incubating our engineered J76-CD8 cryptic-specific TCR cell lines with their respective tetramer coated on a plate and measuring CD69 upregulation after 18 hours. Both cryptic-specific TCRs were able to robustly upregulate CD69, whereas stimulation with an irrelevant antigen or mismatched TCR resulted in background level expression (Fig. 3b).

**Fig. 3.**
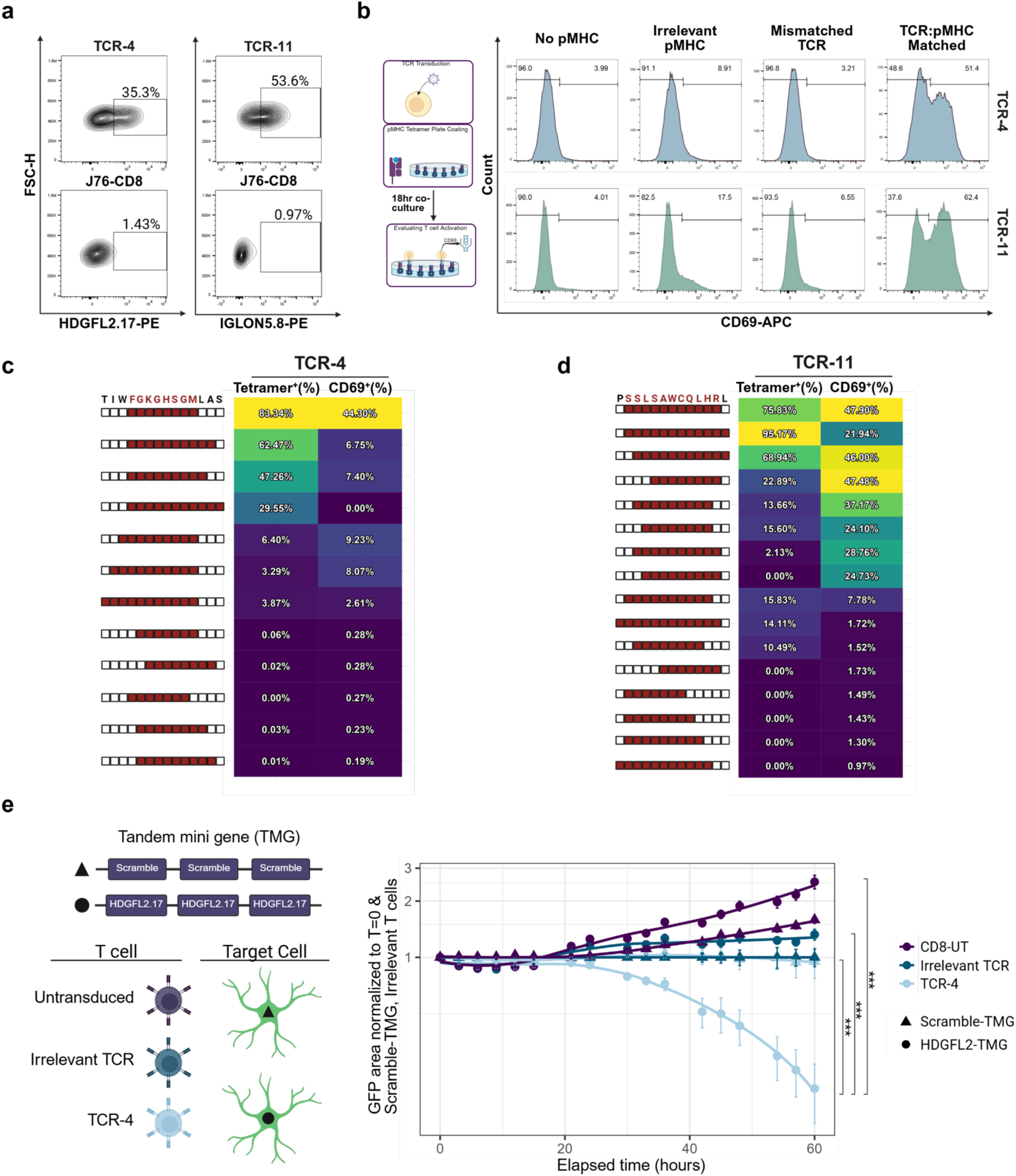
T cells with engineered cryptic epitope specific TCRs can bind and activate in response to cryptic epitopes. **(a)** pMHC tetramer binding of J76-CD8 T cell expressing TCR-4 targeting the HDGFL2 epitope FGKGHSGM, and TCR-11 targeting the IgLON5 epitope SSLSAWCQLHR. **(b)** Flow cytometry histogram plots of CD69 upregulation following 18 hour incubation in a cryptic epitope tetramer coated plate across the following conditions: no pMHC coated on the plate, CMV pp65 (irrelevant) pMHC coated on the plate, irrelevant TCR co-cultured with the matched cryptic epitope pMHC, matched TCR co-cultured with the matched cryptic epitope pMHC. Results for TCR-4 (top row) and TCR-11 (bottom row) are shown. **(c/d)** Evaluation of pMHC tetramer binding and activation of J76-CD8 T cell expressing TCR-4 (c) and TCR-11 (d) across various possible epitope variations of identified cognate cryptic epitopes. The values shown on the heatmaps are background subtracted based on an irrelevant CMV pp65-specific TCR. **(e)** Constructs carrying HDGFL2.17 epitope or scramble epitope were transduced into CCF-STTG1-GFP astrocytes and co-incubated with primary CD8**^+^** T cells transduced to express TCR-4 or irrelevant TCR or untransduced primary CD8**^+^** T cells. GFP area over time was detected as a measure of cell death over time. GFP signals were normalized to the scramble construct with the irrelevant T cell condition. The Kolmogorov–Smirnov (two-sided) test was performed on possible differences in the means of the curves. The standard error for each normalized value is shown across four technical replicates.

To investigate whether TCR-4 and TCR-11 can bind to proteomically processed variants of the originally identified cryptic epitopes, we generated peptides that range from 7-12 aa, each of which contain some portion of the discovered epitope (Fig. 3c,d). Tetramer binding and activation were quite restrictive for TCR-4, and was largely limited to the discovered core epitope, FGKGHSGM.

Extending the peptide at the C-terminus maintained the binding, but significantly diminished CD69 expression. Losing either the phenylalanine or methionine completely abrogated binding and activation (Fig. 3c). On the other hand, TCR-11 proved to tolerate changes to the epitope. The original 11 aa epitope, SSLSAWCQLHR could be reduced to the 8 aa, SAWCQLHR, and maintain the same level of cell activation, with a moderate reduction in binding. Additionally, binding could be enhanced by extending the epitope by one amino acid at the C-terminus, though leading to slightly reduced activation (Fig. 3d).

To further validate specificity and functional capacity of cryptic epitope mediated T cell responses, we transduced an astrocytic-like cell line CCF-STTG1 that is HLA-B*08:01^+^ to constitutively express tagGFP2 (CCF-STTG1-GFP) in order to assess cell survival by monitoring the GFP signal overtime when co-culturing with human primary CD8^+^ T cells transduced to express TCR-4. After co-culturing the CCF-STTG1-GFP engineered to express either the HDGFL2.17 cryptic epitope or a scramble control with TCR-4 transduced primary CD8^+^ T cells, we observed TCR-specific target cell death as measured by GFP loss over time only in the condition where TCR-4 and HDGFL2.17 epitopes were matched (Fig 3e). These data demonstrate that if present, the HDGFL2.17 epitope can be loaded by the cell onto MHC and presented to T cells where they can become targets for killing through TCR recognition.

### Cryptic epitope specific TCR engineered CD8^+^ T cells can kill TDP-43 deficient astrocytes

Given the success of TCR antigen validation using defined epitopes, we wanted to further evaluate whether primary T cells transduced with our TCRs can have a functional response in a TDP-43 proteinopathy relevant biological model. We continued to use the CCF-STTG1-GFP astrocytic-like cell line, as antigen target bearing cells, as glial dysfunction has been found to be relevant in disease pathology^26–30^, and this cell line carries the HLA-B*08:01 allele which is what TCR-4 is restricted to. We used siRNA to knock down TDP-43 in CCF-STTG1-GFP and confirmed that *HDGFL2* cryptic exon is induced by TDP-43 loss in this line (Fig. 4a). This confirms that *HDGFL2* cryptic exon is consistent across cell lines and tissues upon TDP-43 loss.

**Fig. 4.**
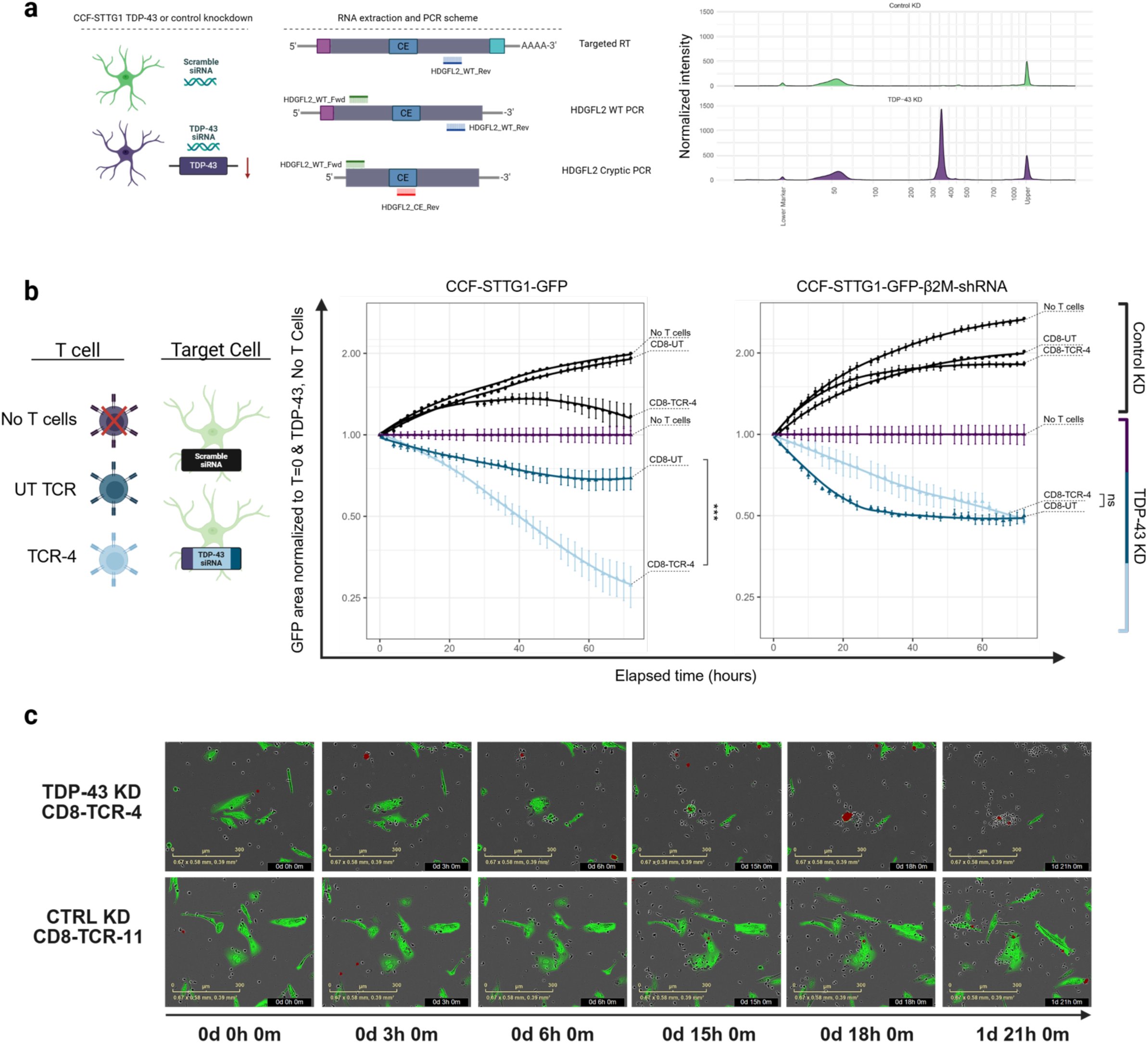
HDGFL2 cryptic epitope specific CD8^+^ T cells can efficiently kill TDP-43 deficient astrocytes. **(a)** siRNA knockdown and HDGFL2 cryptic exon PCR strategy diagram (left) and Tapestation trace of final HDGFL2 cryptic exon PCR product across control and TDP-43 knockdown (right). **(b)** Cytotoxicity of HDGFL2 cryptic epitope specific CD8^+^ T cells toward astrocyte-like cell line, CCF-STTG1-GFP, and its HLA-negative version, CCF-STTG1-GFP-β2M-shRNA, across time as measured by GFP expression. Three T cell conditions across control and siRNA TDP-43 knockdown were measured. CD8-TCR-4: human primary activated CD8^+^ T cells transduced to express TCR-4 and enriched via flow cytometry to be 100% tetramer positive using HDGFL2.17 pMHC tetramer; CD8-UT: primary activated and TCR untransduced CD8^+^ T cells from matching donor; No T cell: no T cells were added to the co-culture. GFP signals were normalized to the No T cell condition. The Kolmogorov–Smirnov (two-sided) test was performed on possible differences in the means of the curves between CD8-TCR-4 and CD8-UT in both panels. ***p = 8.305e-07. ns, p = 0.5041. **(c)** representative images of the CCF-STTG1-GFP:CD8-TCR-4 co-culture across TDP-43 (top) and control (bottom) siRNA knockdown. The red mask marks cells undergoing cell death as indicated by caspase 3/7 dye. The standard error for each normalized value is shown across three technical replicates.

The astrocytic-like cell line CCF-STTG1-GFP treatment with control siRNA resulted in no or limited cell death over time, regardless of the T cell co-culture condition (Fig. 4b, black lines), while TDP-43 siRNA treatment resulted in slightly reduced health comparatively (Fig. 4b, purple lines). Stimulated but TCR untransduced CD8^+^ T cells were able to induce some cell death (Fig. 4b, dark blue line). Strikingly, stimulated CD8^+^ cells transduced to express TCR-4 resulted in a significantly higher amount of cell death (Fig. 4b,c). Using a shRNA targeting β2M to downregulate the expression of MHC class I molecule in CCF-STTG1-GFP line significantly reduced cell death (Fig. 4b, right panel) confirming this TCR-mediated effect and highlighting the TCR specificity toward the cryptic peptide generated by TDP-43 downregulation.

Overall, our result confirms that TDP-43 cryptic peptides could be processed, loaded to MHC class I molecules, and presented on the surface of neural cells, which is in turn recognized by cryptic-epitope specific TCRs. It also signifies that neural cells are susceptible to CD8^+^ T cell mediated killing, and further implicates CD8^+^ T cells in the pathogenesis of TDP-43 proteinopathies.

## Discussion

ALS and IBM are both disorders where TDP-43 proteinopathy represents the predominant pathology^2^. Recently, highly differentiated and clonal CD8^+^ T cells have been observed in ALS and IBM^9,11^. However, their antigen targets and their potential role in ALS and IBM have remained elusive. TDP-43 loss of function leads to the generation of cryptic peptides, which are foreign to the human immune system as the encoding cryptic exons do not exist in the human thymus, raising the possibility that T cells recognizing these peptides are retained. In this study, we highlight significant disease specific expression of HDGFL2 cryptic peptide in IBM skeletal muscle tissue co-localized with TDP-43 nuclear loss using IHC in addition to an increase of T cell infiltrates and MHC class I antigen presentation pathways through RNA sequencing and proteomics. We further identify clonally expanded and highly differentiated CD8^+^ T cell clones which recognize epitopes derived from these cryptic peptides using a high-throughput and multi-dimensional integrated single T cell profiling platform we previously developed^18,19^. Finally, we show that activated CD8^+^ T cells have the capacity to kill astrocytes with TDP-43 loss through cryptic-specific TCR recognition.

Recent work has highlighted clonal and activated T cell populations infiltrating disease-relevant tissue sites across various neurodegenerative disorders^9,10,14,15,24,25,31–37^ and auto-immune diseases of the nervous system^38–42^. However, in many cases, the antigens driving the T cell responses in neurodegenerative diseases are unknown, with one of the main exceptions being PD. It has been shown that aggregates of α-synuclein, the primary pathological feature in PD, can be processed and presented to CD4^+^ and CD8^+^ T cells as self-antigens^34^, that the α-synuclein TCR repertoires of PD patients are highly clonal and diverse^36^ and are present in early stages of disease^35,37^, and associated T cells are in distinctive functional subsets^25^. However, not all aggregation-prone neuronal proteins seem to have the capacity to induce disease-specific T cell activation. T cell responses to amyloid precursor protein, amyloid beta, tau, α-synuclein, and TDP-43 were measured in AD patients, but no difference was observed compared to healthy controls^14^. Similarly, T cell responses to TDP-43 peptides in ALS patients proved to be weak, with healthy controls having moderately higher levels of activation^13^, leaving a knowledge gap on whether TDP-43 pathology and T cell activation are linked.

Differently from other proteins that accumulate in neurodegenerative disease, TDP-43 mislocalisation induces the expression of multiple novel cryptic peptides which are never present in the normal state. We hypothesized these peptides can serve as neo-antigens and our data identify a T cell response against these cryptic peptides, particularly those deriving from IgLON5 and HDGFL2.

IgLON5 is a cell adhesion molecule belonging to the Ig superfamily that is primarily expressed in the brain and testis with poorly understood function^43^. The IgLON5 protein has been recently implicated as a target in an autoimmune and neurodegenerative condition that is characterized by autoantibodies against the IgLON5 protein that is concurrent with neuronal tau deposits^43^. Additionally, a case of a patient with co-existence of neuronal tau pathology along with microglia and neuronal TDP-43 pathology with autoantibodies against IgLON5 has been reported^44^.

HDGFL2 is a ubiquitously expressed histone-binding protein. Cryptic exons derived from HDGFL2 are reproducibly detected via transcript analysis and proteomics in various cell lines including human iPSC-derived neurons, HeLa cells, in addition to CSF samples from patients with ALS/FTD and AD^5,6,20^. Furthermore, two groups have developed antibodies that bind to the cryptic peptide within HDGFL2^6,20^ and showed that HDGFL2 cryptic peptide can be detected in the presymptomatic ALS/FTD^6^ and is significantly increased in brain regions with TDP-43 pathology in FTLD-TDP and AD-TDP, compared to non-TDP-43 controls^20^. In accordance with previous work^7^, we find a florid expression of cryptic exons in IBM muscle tissue, and further we show that HDGFL2 cryptic peptide is also identified in IBM tissue using one of the HDGFL2 cryptic peptide specific antibodies^20^. Intriguingly, we find the HDGFL2 cryptic peptide to co-aggregate in IBM diseased muscle fibers raising the possibility it could further contribute to the aggregation pathogenic cascade. The consistent and widespread detection of HDGFL2 cryptic peptide in CSF, plasma, brain, and muscle supports its role as a potentially strong target for an adaptive immune response. Our findings show that it can induce T-cell mediated cell death and support a potential active role in the degenerative process.

From our single cell RNA and surface protein phenotyping data of the cryptic Tetramer^+^ T cells, we observed an enrichment for a CD8^+^ T_EMRA_ subset in both ALS and IBM conditions, which has previously been reported in ALS-4^9^ and AD^24^. Within this subset, we noted several distinct subpopulations, including a GNLY^Hi^ subset expressing *GZMH*, *GZMB*, and *PRF1*. To investigate whether T cells isolated from other TDP-43 related conditions had shared features with the cryptic-epitope specific cells identified in our study, we identified publicly available single cell TCR sequencing data from the CSF and/or PBMC of AD, PD, ALS, and healthy donors across two distinct datasets^9,24^ (Extended Data Fig. 10a). We identified two TCRs from two different datasets that have nearly identical TCR sequences, one isolated from the PBMCs of an individual with ALS-4 and another from the CSF of an individual with MCI (Extended Data Fig. 10b), compared to two TCRs we identified in this study. This supports the fact that cryptic-epitope specific T cells may have the potential to traffic into the CSF, and that conserved clonotypes can be identified between different individuals. Furthermore, this highlights that our dataset can be utilized as a resource to evaluate potential cryptic epitope specific T cells in already existing or future TCR sequencing datasets.

Modulating adaptive immunity may have the potential to improve clinical outcomes in TDP-43 proteinopathies. It has been shown that individuals with ALS which have a high frequency of activated regulatory T cells (T_regs_) have better survival^10^. Similarly, a recent phase 2b clinical trial utilizing low dose IL-2 (MIROCALS, NCT03039673) has reported that for individuals with low to moderate levels of CSF phosphorylated neurofilament heavy chain (CSF-pNFH, biomarker of disease stage), there was a significant decrease in the risk of death of 48%^45^. Our findings raise the possibility that the effectiveness of this approach is mediated by dampening the response to cryptic peptides and subsequent trials will allow to specifically address this.

Other immunomodulatory clinical trials are ongoing in both IBM and ALS. A phase II/III clinical trial is evaluating the use of an anti-KLRG1 antibody (Ulviprubart) to deplete cytotoxic T cells in IBM (NCT05721573). A Phase 2A trial, the use of tegoprubart targeting CD40-CD40L interaction has shown a decrease in inflammatory markers in people with ALS^46^. Recent trials are also evaluating T_reg_ based cell therapy products. The RAPA-501 (NCT06169176) phase II/III trial is evaluating the use of autologous hybrid T_reg_/TH2 cells manufactured *ex vivo* in high-risk ALS patients. The CK0803 trial (NCT05695521) is evaluating the use of allogeneic, umbilical cord blood derived T_regs_ that express neurotropic homing markers. These approaches, which hope to broadly control the hyperinflammation that is associated with ALS and IBM, could be further refined with an antigen-specific approach in the future. A recent study has shown the neuroprotective value of antigen-specific T cells in a mouse model of CNS injury^47^. This result underscores the value of engineering cells toward a known antigen target. Our work highlights cryptic epitopes as critical T cell antigens and demonstrates the feasibility of engineering T cells to recognize and respond to these antigens.

Altogether, our data demonstrate a previously unknown pathological arm in TDP-43 proteinopathies where the adaptive immune system can be activated through MHC class I antigen presentation pathway, initiating from TDP-43 loss and support the rationale for immune-targeted therapies for TDP-43 proteinopathies.

## Acknowledgements

We thank members of Jiang lab and Fratta lab for stimulating discussions. We thank P. Parikh and S. Salmaso for help with international shipping. We thank NIH Tetramer Core Facility at Emory University for providing MHC monomers. We thank M. Heemskerk lab from Leiden University Medical Center in the Netherlands for the J76 TCR^-^ Jurkat cell line and Paul Thomas lab at St. Jude Children’s Research Hospital for providing human CD8 chain constructs. We are particularly grateful to Penn Center for Cellular Immunotherapies and X. Huang at Drexel University for Incucyte access. We appreciate the statistical analysis guidance provided to us by J. Jin at the Department of Biostatistics, Epidemiology and Informatics of Penn Perelman School of Medicine. A special thank you to R. Balderas at BD Biosciences for providing BD Rhapsody scanner and some of the reagents. This study was supported by researchers at the National Institute for Health and Care Research University College London Hospitals Biomedical Research Centre.

## Funding

This research was supported by

The Chan Zuckerberg Initiative Neurodegeneration Challenge Network Ben Barres Early Career Acceleration Awards 2021-005904 (NJ)

Penn Institute for Immunology and Immune Health pilot award 2022 (NJ)

The Chan Zuckerberg Initiative Collaborative Pairs Pilot Project Awards 2024-338587 (PF, NJ)

NIH R33CA256086 (NJ)

National Institute of Allergy and Infectious Diseases predoctoral training grant T32AI055428 (SC)

MZ is supported by Neuro Research Trust

UK Medical Research Council Senior Clinical Fellowship and MNDA (MR/M008606/1 and MR/S006508/1) (PF)

NIH U54NS123743 (PF)

This research was supported in part by the Intramural Research Program of the NIH, National Institute on Aging (NIA), National Institutes of Health, Department of Health and Human Services; Project number ZIAAG000535.

## Author contributions

Conceptualization: NJ, PF

Data curation: SC, MZ, NJ, PF

Formal analysis: SC, MZ, SK

Funding acquisition: NJ, PF

Investigation: SC, MZ, SK, VS, ALB, VT, LZ, IS, GMM, JMCM, DD, SB, AG, FP, PT, GM, ZL, AQ, LP, MR, GV, EB, MW, AMa, PMM, GS, PF, NJ

Methodology: SC, MZ, NJ, PF

Project administration: SC, MZ, NJ, PF

Resources: AQ, LP, MH, MR, GV, EB, MW, AMa, PMM, GS, PF, NJ

Software: SC, MZ, SK

Supervision: NJ, PF

Writing: SC, MZ, NJ, PF

Writing, review and editing: all authors

## Competing interests

The University of Pennsylvania has applied for patents on the antigens and TCRs.

---

**Extended Data Fig 1.**
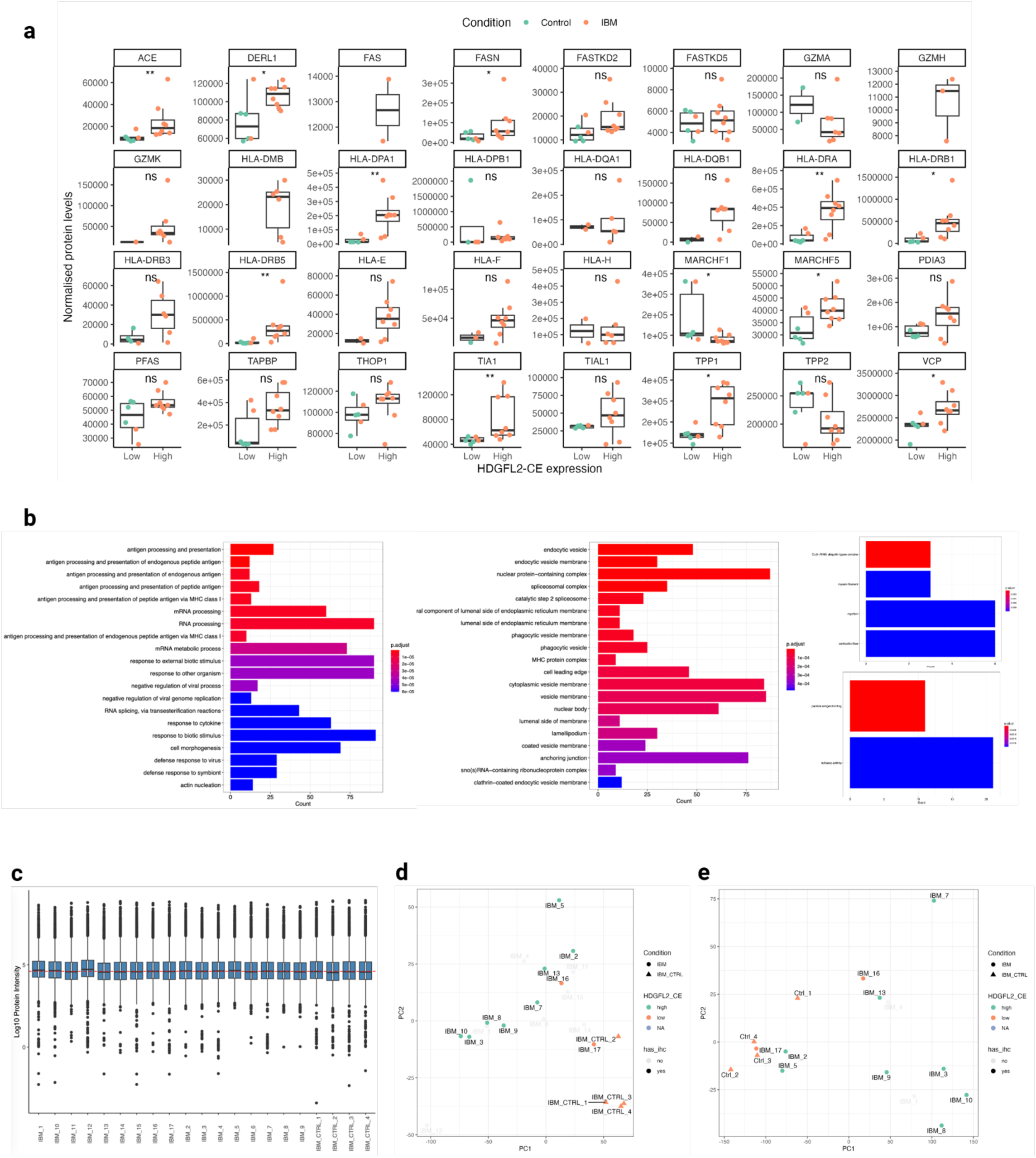
Additional data from the tissue proteomics. **(a)** MHC-I and -II antigen processing pathway expression in controls and IBM cases, divided by condition and coloured based on the upper or lower 50% by TDP43 levels. **(b)** Normalized count distribution for the proteomics cohort. **(c)** GO analysis summary from the proteomics analysis. **(d)** PCA plot from normalised protein expression from the proteomics experiment. **(e)** PCA for the NHNN RNA-seq cohort. Note how the PCA from proteomics and that from RNA are fairly overlapping.

**Extended Data Fig 2.**
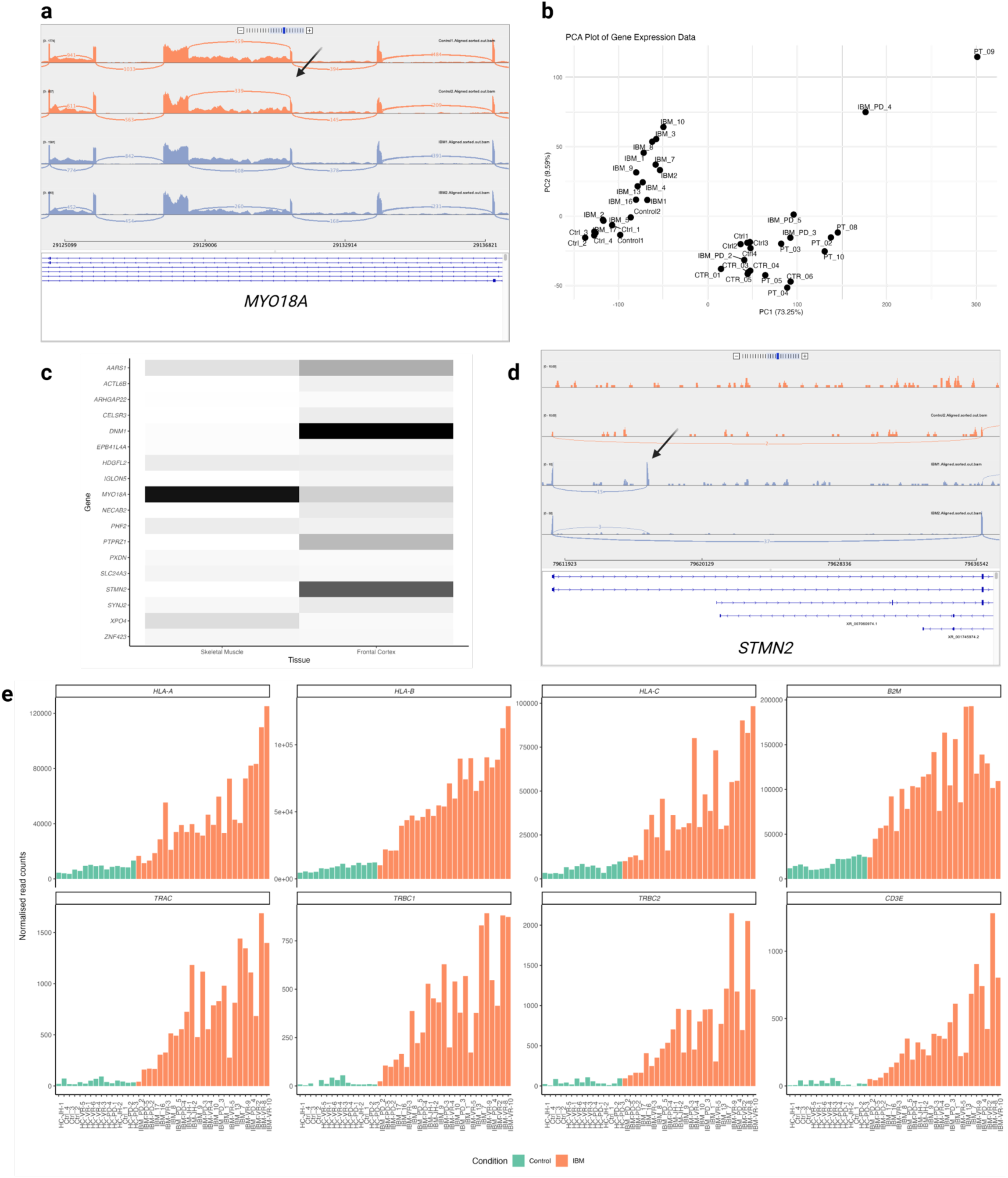
Additional data from RNA-seq analysis. **(a)**. IGV tracks for MYO18A cryptic exon in a subset of controls and IBM muscle biopsy RNA-seq. **(b)** PCA for the full RNA-seq cohort. The clear split between two groups in PC1 is likely driven by different library preparation methods. **(c)** Heatmap showing gene expression of genes with cryptic peptides in skeletal muscle and frontal cortex. Expression data obtained from ASCOT^21^ **(d)** IGV tracks for STMN2 cryptic exon in a subset of controls and IBM muscle biopsy RNA-seq. **(e)** RNA-seq normalized read counts for TCR constant chains (*TRAC*, *TRBC1*, and *TRBC2*), *CD3E*, *HLA-A*, *HLA-B*, *HLA-C*, and beta2-microglobulin (*B2M*) in control and IBM cases.

**Supplementary Table 1.** Tissue analysis cohort.

| **Sample Name** | **Experiment** | **Disease** | **Cohort** | **Alias** |
| --- | --- | --- | --- | --- |
| IBM-2 | Proteomics/IHC/RNA-seq | IBM | UCL_proteomics | IBM-2 |
| IBM-3 | Proteomics/IHC/RNA-seq | IBM | UCL_proteomics | IBM-3 |
| IBM-5 | Proteomics/IHC/RNA-seq | IBM | UCL_proteomics | IBM-5 |
| IBM-7 | Proteomics/IHC/RNA-seq | IBM | UCL_proteomics | IBM-7 |
| IBM-8 | Proteomics/IHC/RNA-seq | IBM | UCL_proteomics | IBM-8 |
| IBM-9 | Proteomics/IHC/RNA-seq | IBM | UCL_proteomics | IBM-9 |
| IBM-10 | Proteomics/IHC/RNA-seq | IBM | UCL_proteomics | IBM-10 |
| IBM-13 | Proteomics/IHC/RNA-seq | IBM | UCL_proteomics | IBM-13 |
| IBM-16 | Proteomics/IHC/RNA-seq | IBM | UCL_proteomics | IBM-16 |
| IBM-17 | Proteomics/IHC/RNA-seq | IBM | UCL_proteomics | IBM-17 |
| Ctrl-1 | Proteomics/IHC/RNA-seq | HC | UCL_proteomics | Ctrl-1 |
| Ctrl-2 | Proteomics/IHC/RNA-seq | HC | UCL_proteomics | Ctrl-2 |
| Ctrl-3 | Proteomics/IHC/RNA-seq | HC | UCL_proteomics | Ctrl-3 |
| Ctrl-4 | Proteomics/IHC/RNA-seq | HC | UCL_proteomics | Ctrl-4 |
| IBM-1 | Proteomics/RNA-seq | IBM | UCL_proteomics | IBM-1 |
| IBM-4 | Proteomics/RNA-seq | IBM | UCL_proteomics | IBM-4 |
| IBM-6 | Proteomics | IBM | UCL_proteomics | IBM-6 |
| IBM-11 | Proteomics | IBM | UCL_proteomics | IBM-11 |
| IBM-12 | Proteomics | IBM | UCL_proteomics | IBM-12 |
| IBM-14 | Proteomics | IBM | UCL_proteomics | IBM-14 |
| IBM-15 | Proteomics | IBM | UCL_proteomics | IBM-15 |
| Control1 | RNA-seq | HC | JH_RNA | HC-JH-1 |
| Control2 | RNA-seq | HC | JH_RNA | HC-JH-2 |
| IBM1 | RNA-seq | IBM | JH_RNA | IBM-JH-1 |
| IBM2 | RNA-seq | IBM | JH_RNA | IBM-JH-2 |
| Ctrl1 | RNA-seq | HC | UP_RNA | HC-UP-1 |
| Ctrl2 | RNA-seq | HC | UP_RNA | HC-UP-2 |
| Ctrl3 | RNA-seq | HC | UP_RNA | HC-UP-3 |
| Ctrl4 | RNA-seq | HC | UP_RNA | HC-UP-4 |
| IBM_PD_2 | RNA-seq | IBM | UP_RNA | IBM-UP-2 |
| IBM_PD_3 | RNA-seq | IBM | UP_RNA | IBM-UP-3 |
| IBM_PD_4 | RNA-seq | IBM | UP_RNA | IBM-UP-4 |
| IBM_PD_5 | RNA-seq | IBM | UP_RNA | IBM-UP-5 |
| CTR_01 | RNA-seq | HC | TS_RNA | HC-TS-1 |
| CTR_03 | RNA-seq | HC | TS_RNA | HC-TS-3 |
| CTR_04 | RNA-seq | HC | TS_RNA | HC-TS-4 |
| CTR_05 | RNA-seq | HC | TS_RNA | HC-TS-5 |
| CTR_06 | RNA-seq | HC | TS_RNA | HC-TS-6 |
| PT_02 | RNA-seq | IBM | TS_RNA | IBM-TS-2 |
| PT_03 | RNA-seq | IBM | TS_RNA | IBM-TS-3 |
| PT_04 | RNA-seq | IBM | TS_RNA | IBM-TS-4 |
| PT_05 | RNA-seq | IBM | TS_RNA | IBM-TS-5 |
| PT_08 | RNA-seq | IBM | TS_RNA | IBM-TS-8 |
| PT_09 | RNA-seq | IBM | TS_RNA | IBM-TS-9 |
| PT_10 | RNA-seq | IBM | TS_RNA | IBM-TS-10 |

## Methods

### Ethics for patient sample collection, MTA, clinical-demographics

For the TCR analysis, we collected PBMCs from n = 4 ALS cases from UCL, n = 5 ALS cases from University of Padua, n = 4 IBM cases from UCL, and n = 7 healthy controls from the human immunology core at the University of Pennsylvania. All cases gave written informed consent in accordance with the principles outlined in the Declaration of Helsinki.

### iMuscle CRISPRi cell line development from human iPSCs

The WTC11 human induced pluripotent stem cell (iPSC) line was used as the parent cell line. The cultures were kept at 37.5°C, 5% CO2 atmosphere, and approximately 90-95% humidity. Cultures were grown using Essential-8™ media with supplement (Life Technologies; #A1517001) and rock 2 inhibitor Chroman-1 (MedChem Express; #HY-15392) was used to aid in single cell survival during passage. Single cells were seeded at 0.033 M/cm^2^ into 5% Matrigel-coated plates (Corning® Matrigel hESC-Qualified, LDEV free; #354277) A 5% Matrigel coating solution was prepared using cold KnockOut™ DMEM/F-12 (Gibco; #12660012) and chilled pipette tips^49^. The coated vessels were incubated at 37.5°C, 5% CO2 for 1 hour or overnight. Cultures were passed before colony overlap, or when approximately 75% confluent. Routine morphology assessments were performed. Stable cell lines with transcription factors under the conditional Tet-On 3G promoter were engineered to rapidly differentiate iPSCs into disease relevant cell types. The Zim3 dCas9 CRISPR inhibition system was delivered via PiggyBac flanked between 5’ and 3’ ITRs co-delivered with the EF1-α Transposase at 2:1 ratio (3 µg total DNA per 1.5 M iPSCs and 5 µLs of Lipofectamine STEM (Thermofisher; STEM00001).The dCas9 protein was cloned under the chicken beta-actin promoter with co-expression of BFP-NLS which was observed after overnight incubation. The polyclonal population was expanded and sorted using the SH800S Cell Sorter (Sony Biotechnology; Model LE-SH800SZFCPL; San Jose, CA) with a sterile sorting chip and a 100 µM nozzle size. A single cell suspension was obtained followed by dilution, cell counting, and resuspension on Essential-8™ media with 50 nM Chroman-1 and Penicillin-Streptomycin (Gibco; #15070063), to prevent contamination after sorting. The iPSCs were kept at 4°C while in suspension and during sorting. Positive (mApple or BFP) and negative controls (non-transfected iPSCs) were used to set the forward (FSC) and side scatter (SSC) gates, followed by single cell gating, and fluorescent marker gating to identify positive populations for each marker. The top 10% double positive (mApple + BFP) population was gated and sorted into 12 well plates coated with Matrigel and Essential-8™ with Chroman-1, selecting cells within the 104-105 intensity. The pre-sorted population was enriched from 5.2% to 96.9% double positive. After sorting, iPSCs were returned to the incubator and allowed to rest for 3-4 hours or overnight before refreshing media. The rock inhibitor was kept until the colonies reached 4-6 cells, passaged 2-3 times before removing the antibiotics, and tested negative for mycoplasma before moving out of quarantine. To rapidly generate skeletal muscle, we subcloned shRNA targeting the miRNA sequence for Oct4 (GTCCGAGTGTGGTTCTGTA) downstream of MyoD1, as described previously^50^. To enable CRISPR inhibition, the Zim3-dCas9 machinery was delivered via PiggyBac followed by hygromycin selection (1.50 mg/mL for 4-7 days).

### Modeling loss of function of TDP-43 via CRISPR interference

The loss of function system was engineered by combining the Zim3 repressor, dCas9 system, and dual sgRNA delivered via a lentiviral transduction. The dual guides were subcloned by creating inserts containing sgRNAs between a tRNA spacer^51^ to deliver 2 non-targeting guides (control) or 2 guides targeting TDP-43. The sequences were either ordered as a GBlock from IDT or amplified by PCR, using primers with sequences related to the desired sgRNA. For the insert, non-targeting sgRNA-1 (gaggcaaccgctctagcgcg) or non-targeting sgRNA-2 (ggttgaaccgcccccgccga) were subcloned under the mouse U6 (mU6) promoter or the human U6 (hU6) promoter respectively. For efficient knockdown of TDP-43, sgRNA-1 (gggaagtcagccgtgagacc) and sgRNA-2 (gaggactaggcccggtctca) were subcloned downstream of the hU6 promoter or the mU6 promoter, respectively. The hygromycin selection cassette followed by eGFP-NLS under constitutive expression (EF1-α) was used as a selection marker, when appropriate. The lentiviral backbone was modified to increase viral titer by subcloning a WPRE upstream of the 3’ LTR.

### Lentiviral Packaging and sgRNA Delivery via Lentiviral Transduction

The sgRNAs for TDP-43 knockdown and non-targeting controls were packaged into lentiviral particles using in-house lentiviral vectors transfected into HEK cells as described previously^52^. After transfection, viral particles were concentrated using Lenti-X™ Concentrator (TakaraBio; #631232) at a 1:10 ratio in dPBS and aliquoted for storage at -80°C until the start of the experiments. The control and TDP-43 sgRNAs were qualitatively evaluated for viral titer by confocal imaging. Briefly, Zim3-dCas9 iPSCs were seeded in Matrigel-coated 96 well plates at 0.04 M/cm^2^ in half medium volume plus Chroman-1 and spun down at 300 g for even coating. The plate was returned to the incubator for 1-3 hours to allow the iPSCs to attach to the surface. A lentiviral aliquot was thawed out on ice, and serially diluted 10 times. After 24 hours of incubation, the media was refreshed, the rock inhibitor was removed and returned to the incubator. The plate was imaged at 72 hours post transduction and the viral titer was estimated based on the percentage of eGFP positive cells. The transduction schedule was designed to maximize guide delivery and disease modeling per cell type. To deliver sgRNAs to target cells, lentiviral particles containing control or TDP-43-targeting sgRNAs were directly added to the iPSCs while in suspension. The iPSCs were singularized and spun down at 300 g for 5 minutes. The cell pellet was resuspended in Essential-8™ with 50 nM Chroman-1 followed by cell counting. Cell populations with a viability above 90% or more were used for the experiments. Based on the viral titer, spin down the desired number of iPSCs, remove the supernatant, and resuspend the cell pellet directly on the lentiviral particles. Allow to rest at room temperature in the biosafety cabinet for 10 minutes. After incubation, resuspend the iMuscle iPSCs on Day 0 media. On Day -2, the cultures were observed, and fresh media with hygromycin was supplied, as needed.

### Direct differentiation of iPSCs into MyoD-1 iMuscle

After puromycin selection, two stable cell lines harboring MyOD-1-shRNA-OCT4 with Zim3-dCas9 machinery were obtained. To generate human skeletal muscle from iPSCs, a transcription factor cassette containing MyoD-1 and shRNA against Oct-4 was cloned under the Tet-On promoter, as described previously^50^. Briefly, on Day 0, iPSCs were singularized and seeded in Myogenic Induction Media (MIM) composed of DMEM/F12, 1X Sodium pyruvate (Gibco; #11360070), 1X NEAA, 110 µM 2-mercaptoethanol (Gibco; #21985023), 10 µg/mL Insulin (R&D; 11376497001), 50 nM Chroman-1, and 2 µg/mL Doxycycline. On Day 1, fresh warm media with a rock inhibitor was provided. The rock inhibitor was removed on Day 2 and daily media changes were performed until Day 6. At Day 7 the MIM was replenished and supplemented with 3 µM CHIR 99021 (Tocris; #4423) for 48 hours. On Day 10, a complete media change was performed using Maturation Media composed of Neurobasal A (Gibco; #10888022), 1X B27 plus, 1X NEAA, 1X GlutaMAX, 1X CultureONE, 10µg/mL NT-3, 50 ng/mL Sonic Hedgehog (Peprotech; #100-45), 100 ng /mL Agrin (R&D Systems; #6624-AG-050), 50 ng/mL Laminin, 200 µM L-Ascorbic Acid (Thomas Scientific; #C988E92), 10 ng/mL human IGF-1(Peprotech; #100-11), 2 µg/mL Doxycycline. Half media changes were performed three times per week from Day 12 until harvest on Day 14.

### Sample Harvest, RNA Purification, and RNA sequencing

The iPS-derived cells were harvested on Day 14 for RNA analysis. The media was aspirated, and 300 µL of TRI Reagent® were added directly to the culture and harvested by scraping. Samples in TRI Reagent®were stored on -80°C for automated high throughput processing using Direct-zol™-96 MagBead RNA (Zymo Research # R2101) and Kingfisher® Apex Automated Purification Instrument with 96 DW head (Thermofisher; #5400930). Briefly, the RNA extraction was performed following the manufacturer’s protocol, using a magnetic bead purification system, DNAse I treatment, and a final elution volume of 55 µL. The RNA concentration was determined Qubit™ Flex Fluorometer (ThermoFisher; #Q33327), and the RNA HS Assay Kit (Invitrogen #Q32855). Samples were normalized to 1 ng/µL and RNA quality was assessed by bioanalyzer. The New York Genome Center prepared libraries using the KAPA Hyper Stranded RNA kit at a sequencing length of 350 bp and depth of 40 M reads. Differential splicing analysis was performed as described below.

### Publicly available data

Cross-linking and immunoprecipitation (CLIP) data, used for the analysis of the binding of TDP-43 and other RNA-binding proteins around and within CEs, was downloaded from the POSTAR database^48^ and visualised by custom scripts in R. Information about gene expression in different tissues was obtained from ASCOT.^53^ Genes of interest were selected and displayed using the tidyverse package in R.

### Cryptic Exon Epitope Prediction and Synthesis

The pVACbind feature from pvactools^23^ (v3.1.2) was utilized to evaluate epitope binding across the following HLA: HLA-A*01:01, HLA-A*02:01, HLA-A*03:01, HLA-A*24:02, HLA-B*07:02, HLA-B*08:01, HLA-B*35:01, HLA-B*40:01, HLA-C*07:02. For each cryptic exon amino acid peptide sequence, 8 amino acids from the left and right flanking exon were included and 8 – 11mer sliding window epitopes were considered using the following MHC binding algorithms: MHCflurry, NetMHC, NetMHCpan, PickPocket, SMM, and SMMPMBEC. An epitope was considered a binder if any of the 6 algorithms called it a either a top 0.5% binder (HLA-B and HLA-C), or a top 1% binder (HLA-A). This strategy resulted in 379 distinct peptide:MHC pairs across a total of 331 epitopes. All predicted epitopes were synthesized by Genscript.

### High-throughput screening of peptide-MHC candidates for non-specific binding and noise

Our lab has data suggesting that a few short 8 – 11mer amino acid peptides have the capacity to non-specifically bind to streptavidin and subsequently bind to the cell surface, independent of MHC interaction (data unpublished). This phenomenon greatly increases noise and reduces the specificity of Tetramer:TCR binding which is critical, especially in high-throughput screening. Therefore, following the synthesis of all peptides of interest and prior to UV-mediated loading of peptides onto HLA molecules, the peptides were screened for non-specific interactions with APC and PE Streptavidin. In this method, APC or PE Streptavidin is co-incubated with each peptide individually at a final concentration of 30nM and 150uM respectively in the absence of any HLA monomer. The streptavidin-peptide mixture is then co-incubated with either J76 or primary T cells for 30 minutes, then washed and evaluated using flow cytometry. Any peptide exhibiting more than 2% APC or PE signal was flagged and removed from the final tetramer panel.

### High-dimensional, tetramer-associated T cell antigen receptor (TCR) sequencing (TetTCR-SeqHD)

TetTCR-SeqHD^18,19^ is a previously described single cell sequencing method developed in our lab capable of profiling T cell cognate antigen specificities, T cell receptor sequences, gene expression, and surface protein expression. In brief, a large library of fluorescently labeled and DNA barcoded pMHC tetramers are generated through peptide synthesis and ultraviolet light-mediated peptide exchange. Next, pMHC tetramers are pooled and used to stain T cells along with barcoded antibodies (Abseq) for surface phenotyping and sample multiplexing. Antigen-bound T cells are sorted and are loaded onto a BD Rhapsody single cell capture chip. Following several PCR processing steps, the libraries are sequenced with next generation sequencing, and computationally processed to generate cell-UMI matrices across gene expression, surface phenotype, TCR expression, and tetramer-specificity.

### Informatic Processing of Gene Expression, Abseq, Sampletag, TCR, and Tetramer

Raw single cell sequencing files were processed using a combination of BD Rhapsody’s single cell pipeline in addition to a custom pipeline which generates tetramer cell-UMI matrices. Seurat (v4.0.2) was used to generate a merged Seurat object consisting of all experimental runs. RNA and Abseq counts were normalized using the SCTransform function. A weighted nearest neighbor graph was generated using the FindMultiModalNeighbors function utilizing dimensionally reduced RNA and Abseq modalities, and a UMAP was constructed. Clusters were manually annotated using a combination of the differentially expressed genes and Abseq per cluster. Tetramer specificities per cell were assigned using a modified version of the Seurat HTODemux function. Specificities were refined by assigning *clonal specificities* such that at least 60% of the cells in a given clonotype as defined by their CDR3b amino acid sequence must share the same epitope assignment. TCR clonality was visualized with the scRepertoire (v.1.7.2). The frequency of clonotypes was binned by patient with the following parameters: Singleton = 1, Small = 3, Medium = 10, Large = 15, Hyperexpanded = 100, and clones were determined by the CDR3b amino acid sequence.

### TCR Cloning, Transfection, and Transduction

The alpha and beta CDR3 sequences identified through the BD Rhapsody Sequence Analysis pipeline were aligned to the IMGT TCR reference to generate full length TRBV and TRAV sequences. The sequences were codon-optimized and IDT eblocks were ordered for each fragment. Fragments were assembled using HIFI assembly with an optimized backbone compatible with lentiviral transfection. The assemblies were transformed into NEB Stable Competent E.coli (High Efficiency) and a standard cloning protocol was used to generate plasmids. Sanger sequencing validated TCR plasmid were transfected into HEK293T cells in combination with a lentiviral packaging mix to generate virus. Virus was collected and concentrated for subsequent TCR transduction.

### TCR Binding and Activation Validation

The binding specificity of the TCR-transduced cell lines was validated using flow cytometry with pMHC tetramers. To create pMHC complexes, 6 μL of 200 μM peptide and 2 μL of 0.2 mg/mL monomer were combined and UV-exchanged for an hour. After incubating at 4°C overnight, the pMHC complexes were first evaluated for stability using streptavidin-coated beads (6-8 μm) and incubated at 37°C for 1 hour. Following washing, PE-labeled anti-human β2m monoclonal antibody was used to stain for 30 minutes at 4°C and then analyzed by flow cytometry. For TCR binding validation, the stable pMHC complexes were tetramerized with 4 μL of 0.045 mg/mL fluorescently-labeled PE-streptavidin. 50,000 transduced cells were used for each condition and stained with 1.5 μL of tetramer in 50 μL of FACS buffer (PBS + 2% FBS + 5mM EDTA) for 1 hour at 4°C before washing and running on flow cytometry. Untransduced J76-CD8 cell lines and irrelevant peptides were used as negative controls. To evaluate the activation of the TCR-transduced cell lines by the cognate peptides, we first coated 0.1 μg of streptavidin in an ELISA 96-well plate and incubated at room temperature overnight. After washing with 0.05% Tween20 in PBS, blocking buffer (2% BSA in PBS) was added and incubated at room temperature for 30 minutes. The pMHC complexes were diluted to 450 ng with blocking buffer for each reaction. CMV-specific pp65 pMHC with the corresponding transduced cell lines were used as controls for the activation assay. 100 μL of diluted pMHC monomer or 500 μM biotin solution was added to each sample or control wells, respectively, followed by incubation at 37°C for 1 hour. 125,000 TCR-transduced cells were added and incubated at 37°C for 18 hours. After washing with FACS buffer, the cells were stained with APC-CD69 antibody for 30 minutes and evaluated using flow cytometry.

### CCF-STTG1 TDP-43 Knockdown: RNA-seq and HDGFL2 cryptic exon PCR Analysis

CCF-STTG1 cells were seeded in twelve-well plates at a density of 150,000 cells / well in 800 μL of RPMI 1640 +10% FBS without antibiotics. The following day, cells were transfected with either MISSION siRNA targeting human TARDBP (cat. EHU109221) or with MISSION siRNA Universal Negative Control #1 (cat. EHU109221). siRNA-Lipofectamine complexes were generated by combining 40 pmol of siRNA in 100 μL of OptiMem with 2 μL of Lipofectamine 2000 in 100 μL of OptiMem. After a 20 minute incubation, the combined mixture was added to the cells. Following a 48 hour incubation, RNA was isolated from each well by following Zymo’s Quick-RNA Miniprep Kit (cat. R1054). RNAseq libraries were prepared using NEBNext Ultra II Directional with Poly-A selection and sequenced using Novaseq X 2x150, with 40M PE reads per sample.

To evaluate the HDGFL2 cryptic exon expression, a modified strategy described by Irwin et al.^6^ was implemented. Briefly, reverse transcription was performed with a reverse primer targeting the exon region which follows the cryptic exon inclusion of HDGFL2: 5’-CTCGGAGGCTTCTTCACAGACA-3’. PCR was then performed using a primer pair targeting the exons flanking the cryptic exon inclusion: Fwd-primer 5’-AAGACGCCTGCGCTAAAGAT-3’, Rev-primer 5’-CTCGGAGGCTTCTTCACAGACA-3’. Finally, HDGFL2 cryptic exon expression was evaluated with the following primer pair: Forward primer 5’-AAGACGCCTGCGCTAAAGAT-3’ and HDGFL2 CE Reverse primer 5’-GCTTCCCTCCCTTCTGATGC-3’.

### CCF-STTG1:T cell Co-culture

The CCF-STTG1-GFP cell line was first generated by virally transducing CCF-STTG1 cells to constitutively express tagGFP2 using the viral transduction strategy described above. These cells were then plated at a seeding density of 2,000 cells/well in a flat 96-well plate in 50 μL of RPMI+10% FBS without antibiotics. The following day, TDP-43 or control siRNA was complexed with Lipofectamine 2000 as described above, and 1 pmol of siRNA was added to the cell culture. 24 hours following the siRNA transfection, cells were thoroughly washed with media and then co-incubated with the following primary T cell conditions: CD4/CD8 transduced, CD4/CD8 untransduced, and CD4/CD8 naive. In each condition, CD4^+^ or CD8^+^ T cells were enriched using EasySep Human CD4^+^ or CD8^+^ negative enrichment kit. The transduced and untransduced cells were both initially stimulated with ImmunoCult Human CD3/CD28 T Cell Activator. The naive cell conditions were unstimulated and CD45RA+CCR7+ cells were FACS sorted. A portion of the T cells were lysed and RNA was collected for RNAseq prior to co-incubation. The T cells were resuspended in RPMI + 10% FBS + 1x Caspase-3/7 Red and 5,000 T cells / well were co-incubated with the siRNA-treated CCF-STTG1-GFP cells. Each condition was performed with 3-4 replicates. Cells were monitored using the IncuCyte system where scans were taken every 3 hours.

### CCF-STTG1 B2M-deficient shRNA generation

To generate stable B2M deficient shRNA cell lines, the pLKO.1 shRNA backbone was used to target the following sequences: 5’-AGTTAAGCGTGCATAAGTTAACTCGAGTTAACTTATGCACGCTTAACT-3’ and 5’-TTCAATCTCTTGCACTCAAAGCTCGAGCTTTGAGTGCAAGAGATTGAA-3’. The shRNA assemblies were transformed into NEB Stable Competent E. coli (High Efficiency), and standard molecular cloning protocols were used to generate plasmids, which were validated by full plasmid sequencing. The validated plasmids were transfected with a lentiviral packaging mix into HEK293T cells to produce lentiviral particles, which were collected, filtered, and concentrated. The virus was subsequently used to transduce CCF-STTG1-GFP astrocyte cells to establish stable shRNA-expressing cell lines.

### GLIPH2 TCR analysis

We performed GLIPH2^54^ clustering using our experimentally identified TCRs along with reference datasets from literature^9,24^. We further filtered the output from GLIPH2 to identify TCRs with a higher probability of sharing specificities. The GLIPH2 output was filtered to include only patterns that contained samples from both our experimental dataset and our reference set from the literature. For each pattern, we calculated the minimum Levenshtein distances between the TCRβ CDR3 sequences of the two groups as well as between TCRα CDR3a sequences, excluding any empty or missing values. The TRBV allele was compared to identify matching V gene segments between the groups within each pattern. We included patterns which contained CDR3 Levenshtein distances of one or less along with a matching TRBV allele between our experimental dataset and the reference.

### Muscle biopsies

The use of samples for the present study was approved by Brain UK. All cases gave written informed consent in accordance with the principles outlined in the Declaration of Helsinki. Anonymisation of study samples was performed by a member of the clinical care team who was not directly involved in the study. We selected cases based on the following inclusion and exclusion criteria:

1. IBM group: Any person diagnosed with IBM by a qualified clinician, according to internationally accepted diagnostic criteria^55^
2. Controls: Normal muscle tissue, from patients investigated for cramps or fatigue with normal examination and neuro-physiology tests and normal histology

Quadriceps muscles from 17 IBM and 4 control cases were biopsied, flash-frozen in liquid nitrogen, and stored at -80 °C in the Neuropathology department of the National Hospital for Neurology and Neurosurgery (NHNN) in London, UK.

### Immunohistochemistry

For IHC, 8 𝛍m sections from 10 IBM cases and 4 controls were cut with a cryostat. DAB-staining consisted of 10 minutes of fixation with 4% PFA, followed by blocking endogenous peroxidases with 3% H2O2 in methanol. Pre-blocking in 10% milk/TBS-T for 30 minutes was followed by a 2-hour incubation with the following primary antibodies:

- TDP-43, mouse monoclonal (Abnova, 1:800)
- phosphoTDP-43, mouse polyclonal (Cosmo, 1:4000)
- p62, mouse monoclonal (Abcam, 1:100)
- HDGFL2 cryptic peptide, rabbit polyclonal (1:500)^20^

This was followed by a 30-minute incubation with secondary antibodies. The visualization step consisted of 30 minutes of incubation with ABC complex and DAB staining for 3 minutes, followed by counterstaining in Mayer’s hematoxylin and mounting overnight prior to visualization on the microscope and scanning. Quantification of immune infiltration via IHC was performed by visually assessing the extent of immune cells in the whole sections in a blind way, and confirmed by a neuropathologist. Quantification of TDP-43, p62 inclusions, and HDGFL2 cryptic peptide burden analyses were based on DAB staining signal and performed in a blind, semiquantitative fashion, and confirmed by a neuropathologist.

### Proteomics

4 25 𝛍m thick flash-frozen muscle biopsies from each IBM and control case were collected and 20 mg of tissue were dissolved in 100 𝛍l of lysis buffer, consisting of 50 mM Tris-HCl pH 8, 50 mM NaCl, 1% v/v SDS, 1% v/v Triton X-100, 1% v/v NP-40, 1% v/v Tween 20, 1% v/v glycerol, 1% sodium deoxycholate (w/v), 5 mM EDTA pH 8, 5 mM dithiothreitol, 5 KU benzonase and protease inhibitors. Following homogenization and spinning at 20000g at 4 °C for 15 minutes, cells were snap-frozen.

The sample preparation for mass spectrometry (MS)-based proteomics was performed on our previously published fully automated pipeline^56^. Briefly, Cell lysates were incubated at 65 °C for 30 min, followed by addition of 10 mM iodoacetamide and 30 min incubation in the dark. The denatured proteins were measured and normalized using an automated colorimetric assay on a Bravo (Agilent Technologies) robot. The protein enrichment and tryptic on-beads digestion were carried out on a KingFisher robot. The final resulting tryptic peptides were separated on a nano column (75 μm × 500 mm, 2 μm C18 particle) using a 2-h efficient linear gradient (Phase B, 2-35% ACN) on an UltiMate 3000 nano-HPLC system. We used data independent acquisition (DIA) discovery proteomics on a hybrid Orbitrap Eclipse mass spectrometer. Specifically, MS1 resolution was set to 120K, and MS2 resolution was set to 30K. For DIA isolation, the precursor range was set to 400–1000 m/z, and the isolation window was 8 m/z, resulting in 75 windows for each scan cycle (3 s). High collision dissociation was used for fragmentation with 30% collision energy. The AGC target was set to 800% for MS2 scan (improve MS2 spectra quality). The MS2 scan range was defined as 145–1450 m/z, which covers most fragment ions. The database search was performed at the protein level by Spectronaut software 16.0. We used a direct DIA, which does not require a preconstructed high-quality project-specific spectral library but only uses a proteome sequence file. Specifically, the MS/MS spectra were searched against a proteome reference database containing 17,125 proteins (reviewed) mapped to the human reference genome (UP000005640) obtained via the UniProt Consortium. All spectra were allowed ‘‘trypsin and lysC’’ enzyme specificity, with up to two missed cleavages. Allowed fixed modifications included carbamidomethylation of cysteine and the allowed variable modifications for whole proteome datasets were acetylation of protein N-termini, oxidized methionine. The false discovery rates of peptide and protein were all set as 1%. The post-analysis and data visualization were carried out using Protpipe informatics suite^57^. Custom scripts written in R software were used to analyze and visualize the normalised count data.

### RNA-seq analysis

RNA-seq data for n = 2 controls and n = 2 IBM were publicly available and were obtained from the authors (Johns Hopkins - JH cohort)^7^. Novel data included a cohort from Padua (PD), including n = 4 controls and n = 4 IBM, and a cohort from Verona (VR), including n = 5 controls and n = 7 IBM. Moreover, we sequenced 12 IBM and 4 control from our skeletal muscle biopsy cohort (Supplementary Table 1). 4 25 𝛍m thick flash-frozen muscle biopsies from each IBM and control case were collected and 20 𝛍g of tissue were dissolved in QIAzol Lysis Reagent (QIAGEN) prior to RNA extraction using the miRNeasy mini kit (QIAGeN). For the PD cohort, sequencing libraries were prepared with polyA enrichment and sequenced at UCL Genomics with the following specifics: 2×150 bp, depth >40M/sample. For the VR cohort, sequencing libraries were prepared with polyA enrichment and sequenced by Novogene with the following specifics: 2x150 bp, depth >40M/sample. For the NHNN cohort, sequencing libraries were prepared with ribo-depletion and sequenced by Novogene with the following specifics: 2x150 bp, depth >60M/sample. Raw sequences in fastq format were trimmed using Fastp^58^, and aligned using STAR (v2.7.0f)^59^ to the GRCh38 reference genome with gene models from GENCODE v42^60^, including the generation of splice junction counts. For visualisation purposes, aligned BAM files were sorted and indexed by the read coordinates location using samtools^61^. To generate normalised gene counts, trimmed fastqc files were aligned to the transcriptome using Salmon (v1.5.1)^62^, before running DeSEQ2^63^ without covariates, using an index built from GENCODE v42^60^. The DESeq2 median of ratios, which controls for both sequencing depth and RNA composition, was used to normalise gene counts. We kept genes with at least 5 counts per million in more than 2 samples. The alignment pipeline is implemented in Snakemake (v5.5.4)^64^, and run on the UCL high-performance cluster.

To assess the presence of cryptic exon encoding for cryptic peptides in the IBM RNA-seq cohort, we used the splice junction output from STAR^59^. A cryptic exon was considered detected in a sample if there was at least one uniquely mapped spliced read supporting either the acceptor or the donor junction, and for quantification purposes, only the higher between the two was considered. Cryptic splice junctions were exported as a BED file that was used as a query to quantify cryptic exon expression in the IBM RNA-seq data. This returned a table with coordinates, gene name, and how many spliced reads in each sample supported that junction. Lastly, we normalised read count by the total number of mapped read per sample. This pipeline uses bedtools^65^ and bash parsing with awk and is implemented in Snakemake v5.5.4^64^. Data visualisation was run on R using custom scripts.

## Notes

### Summary of Updates

We have updated the author list, along with one additional funding source for this revision.

